# Analysis of lifespan across Diversity Outbred mouse studies identifies multiple longevity-associated loci

**DOI:** 10.1101/2024.11.20.624531

**Authors:** Martin N. Mullis, Kevin M. Wright, Anil Raj, Daniel M. Gatti, Peter C. Reifsnyder, Kevin Flurkey, Jonathan R. Archer, Laura Robinson, Andrea Di Francesco, Karen L. Svenson, Ron Korstanje, David E. Harrison, J. Graham Ruby, Gary A. Churchill

## Abstract

Lifespan is an integrative phenotype whose genetic architecture is likely to highlight multiple processes with high impact on health and aging. Here, we conduct a genetic meta-analysis of longevity in Diversity Outbred (DO) mice that includes 2,444 animals from three independently conducted lifespan studies. We identify six loci that contribute significantly to lifespan independently of diet and drug treatment, one of which also influences lifespan in a sex-dependent manner, as well as an additional locus with a diet-specific effect on lifespan. Collectively, these loci explain over half of the estimated heritable variation in lifespan across these studies and provide insight into the genetic architecture of lifespan in DO mice.

## INTRODUCTION

Lifespan is a quantitative phenotype that can summarize health and fitness information across populations and physiological systems. When seeking to discover correlates that are potentially causative, the integrative nature of this phenotype provides both advantages and disadvantages to researchers. Disadvantages relate to the plethora of unrelated contributors: each correlate likely makes only a small contribution to the phenotype, limiting the statistical power to discover it. When discovered, the non-specific nature of the phenotype also confounds interpretation: the correlate could affect any unknown sub-population or subset of health phenomena. On the other hand, those correlates that do emerge are likely to have either the highest and/or broadest impacts on overall health for the population, which provides an advantage in terms of prioritization. These principles apply to any covariate of lifespan, including alleles that drive the genetic analyses described herein.

In human populations, the heritability (*h*^2^) of lifespan is low, consistently reported to be under 20% (e.g. Kaplanis et al. 2018; Van Den Berg et al. 2017) and under 10% when accounting for assortative mating (Ruby et al. 2018). This limits the power of quantitative trait locus (QTL) discovery. Large cohorts have nonetheless empowered such discovery, but the significance of many loci vary across distinct populations and study designs (e.g. Fortney et al. 2015; McDaid et al. 2017; Wright et al. 2019). Outbred mouse populations provide orthogonal resources to study the genetics of mammalian lifespan, with advantages and disadvantages versus human populations. The limited sample sizes of typical mouse experiments versus modern human cohorts are a disadvantage. Advantages include: well-balanced allele frequencies, which increase statistical power to detect loci; lack of substantial population structure, which confounds genotype/phenotype correlations in human populations but has minimal impacts in mouse populations (Churchill et al. 2012; Ghazalpour et al. 2012; Wang et al. 2021); and the ability to enroll mice at a young age and then follow them until the end of their natural lifespans, facilitating prospective analyses of lifespan. Finally, treatments and environmental perturbations may be applied to mouse populations, allowing the effects of these interventions on lifespan and other health-related phenotypes.

Dozens of lifespan-associated QTL have been mapped in mice using a variety of techniques (Hook et al. 2018). These include the use of congenic mouse strains to investigate the potential effects of particular loci (Smith and Walford 1977), with subsequent research utilizing backcrosses (Yunis et al. 1984). Recombinant inbred lines (Peirce et al. 2004) and intercrosses (Yuan et al. 2013) have also been used to identify lifespan QTL, but those study populations contained limited genetic diversity and recombination, resulting in coarse mapping resolution and restricting the ability to identify candidate genes. Advances in genome sequencing and the development of large-scale intercrosses (Miller et al. 2007; Bou Sleiman et al. 2022) and outbred populations (Churchill et al. 2012; Svenson et al. 2012) are now beginning to enable the mapping of lifespan at a higher precision in populations with greater genetic diversity.

Several interventions are known to extend mouse lifespan (Miller et al. 2007) and healthspan, including multiple approaches to dietary restriction (DR) (Weindruch et al. 1986; Anderson, Shanmuganayagam, and Weindruch 2009; Mitchell et al. 2019) and several compounds, including the drug rapamycin (Heitman, Movva, and Hall 1991; Bjedov et al. 2010; Harrison et al. 2009). These interventions are robust across mouse genotypes, as evidenced by their validation in genetically heterogeneous populations (Di Francesco et al. 2023). Despite their robustness, consensus on the physiological mechanisms-of-action for these interventions is lacking (Green, Lamming, and Fontana 2022; Papadopoli et al. 2019). Mechanistic insight could be gained through study of interacting effects between genetic loci and either of these interventions, considered as environment covariates (gene-by-environment interactions; GxE) (Ottman 1996). Genetic loci responsive to DR are mapped in mice for traits related to cardiac physiology (Zhang et al. 2022), but GxE effects on lifespan remain unexplored.

Here, we present data from three previously unpublished lifespan studies conducted using Diversity Outbred (DO) mice (Churchill et al. 2012): the Harrison, Svenson, and Shock studies. We used these data, together with DO lifespan data from the published DRiDO study (Di Francesco et al. 2023), to evaluate the robustness of lifespan-extending interventions and characterize the genetic architecture of DO mouse lifespan. We found dietary restriction and rapamycin treatment to robustly increase lifespan across all relevant studies (Harrison, Svenson and DRiDO). For the three studies that included genotypes (Harrison, Shock and DRiDO), we performed genetic analysis of each individual study, identifying five non-overlapping QTL associated with lifespan. Further meta-analysis across all three genetic studies revealed three additional QTL. Finally, we performed a genome-wide genetics-by-environment (GxE) scan, revealing one locus whose effect on lifespan was modified by dietary intervention.

## RESULTS

Lifespan data analyzed here derived from four separate studies of DO mice, referred to herein as the Harrison study, the Svenson study, the Jackson Laboratory Nathan Shock Center (Shock) study, and the Dietary Restriction in Diversity Outbred mice (DRiDO) study. Three are reported here for the first time: Harrison, Svenson, and Shock. The fourth was previously described (Di Francesco et al. 2023). These studies were all conducted independently and involved different cohorts, facilities, technicians, and methodologies (**Data S1**). Each study also had its own design, investigating dietary, drug, and/or sex effects on lifespan. Those designs are summarized in **Figure 1B-E** and described in more detail in the **Methods**. Genotype data was additionally collected from the Harrison, Shock, and DRiDO studies (see **Methods**).

**Figure 1.**
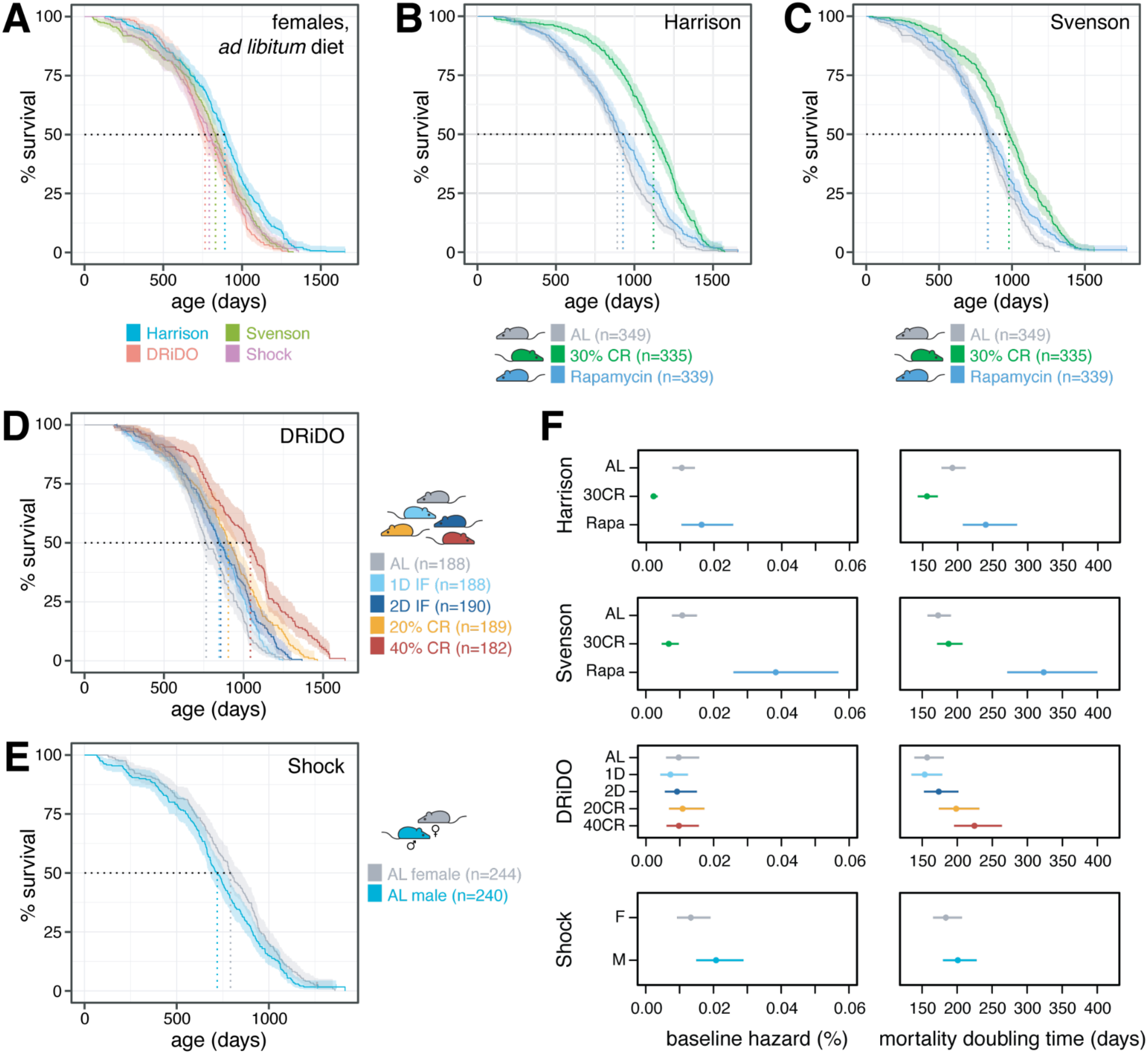
Survival in DO mice across four independent studies. **A,** Kaplan-Meier survival curves for *ad libitum* females in each of the four studies. Dashed vertical lines indicate median survival in days and bands around each curve indicate 95% confidence intervals. Vertical bars on each curve indicate censorship events. **B,** Kaplan-Meier survival curves by intervention in the Harrison study. **C,** Kaplan-Meier survival curves by intervention in the Svenson study. **D,** Kaplan-Meier survival curves by dietary intervention in the DRiDO study. **E,** Kaplan-Meier survival curves by sex in the Shock study. **F,** Baseline hazard and mortality doubling times and 95% confidence intervals estimated by a Gompertz log-linear model.

Despite varied designs, one control group was shared by all four studies: female mice fed an *ad libitum* (‘AL’), nutrient-rich chow diet (**Methods)**. Median lifespans for this category were 794 days (95% CI = 754, 855), 832 days (95% CI = 800, 860), 891 days (95% CI = 863, 922), and 765 days (95% CI = 728, 839) in the Shock, Svenson, Harrison, and DRiDO studies, respectively, and may reflect differences in the design and execution of the studies, including housing, feeding time, and facility (**Figure 1A, Methods**). Lifespan in AL females varied significantly by study (log-rank test, ꭓ^2^ = 33.54, p = 2.48*10^-7^, **Figure 1A**); however not all of the studies were statistically different from one another (**Table S1**), and study explained a significant but small proportion of total variance in lifespan among AL females (ANOVA, PVE = 1.3%, p = 2.9*10^-3^). Relative to the DRiDO study, the Cox hazard ratio was insignificant for the Shock and Svenson studies (0.88 and 0.85, respectively; p = 0.2 and 0.078, respectively), but hazard was significantly reduced in the Harrison study (0.61, p = 2.03*10^-7^). For downstream statistical analyses, study was included as a covariate wherever relevant.

### Rapamycin and caloric restriction both extended lifespan in DO mice

The Harrison and Svenson studies enrolled 1023 and 952 female mice, respectively, and both probed the effects of 30% caloric restriction (‘30CR’) and rapamycin drug intervention (‘Rapa’) on lifespan. Despite beginning treatment at 16 months of age, rapamycin increased lifespan relative to the AL controls in both the Harrison study (**Figure 1B**; log-rank test, ꭓ^2^ = 8.23, p = 4.13*10^-3^) and the Svenson study (**Figure 1C**; log-rank test, ꭓ^2^ = 11.14, p = 8.44*10^-4^). Rapamycin did not increase median lifespan relative to AL controls, possibly because treatment began in midlife. Nonetheless, Cox hazard ratios for rapamycin-treated animals versus AL controls were 0.77 (95% CI = 0.66, 0.90) and 0.74 (95% CI = 0.63, 0.86) in the Harrison and Svenson studies, respectively.

Caloric restriction at 30% also increased lifespan in both the Harrison and Svenson studies. In the Harrison study (**Figure 1B**), median lifespan was increased by 230 days (891 days for AL versus 1121 days for 30CR; a 25.8% increase); lifespan extension was statistically significant (log-rank test, ꭓ^2^ = 96.14, p = 1.07*10^-22^); and mortality hazard was reduced (Cox HR = 0.47, 95% CI = 0.4-0.56). In the Svenson study (**Figure 1C**), median lifespan was increased by 150 days (832 for AL versus 982 days for 30CR; an 18% increase); lifespan extension was again statistically significant (log-rank test, ꭓ^2^ = 75.72, p = 3.28*10^-18^); and mortality hazard was again reduced (Cox HR = 0.52, 95% CI = 0.44, 0.61).

The DRiDO study measured lifespans for 937 female mice randomly assigned to five different diet groups at six months of age: *ad libitum* (‘AL’), one and two consecutive days of intermittent fasting (‘1D’ and ‘2D’, respectively), and 20% and 40% caloric restriction (‘20CR’ and ‘40CR’, respectively). As previously published (Di Francesco et al. 2023) and depicted in **Figure 1D**, all of these variations on dietary restriction significantly increased lifespan.

### Lifespans were longer for female versus male DO mice

The Shock study enrolled 244 female and 240 male mice and did not include any dietary or drug intervention groups: all mice were fed an AL diet similar to the Harrison, Svenson, and DRiDO studies (see **Methods**). The median lifespan of female mice was 75 days longer than male mice (**Figure 1E**; 794 days for females versus 719 days for males; log-rank test, ꭓ^2^ = 5.33, p = 2.09*10^-2^), with mortality hazard greater in males (Cox HR = 1.24, 95% CI = 1.03-1.48). This result was reminiscent of the slight female survival advantage reported for genetically diverse UM-HET3 mice (Cheng et al. 2019; Bou Sleiman et al. 2022).

### The effects of rapamycin and caloric restriction on rates of demographic aging were inconsistent between studies

The mortality doubling time is defined as the rate of increase in mortality hazard with increasing age (Finch, Pike, and Witten 1990), and is expressed here as the period of time across which mortality hazard doubles when lifespans are fit to a Gompertzian log-linear hazard model . It was previously reported that CR slowed demographic aging in the DRiDO study for the 20% and 40% CR groups, without having a significant effect on the baseline hazard (Di Francesco et al. 2023).

We estimated both the rates of demographic aging and baseline hazard for each sex and intervention group across all four studies (**Figure 1F**). Although 30% CR extended lifespan reproducibly in both the Harrison and Svenson studies, the effects on the mortality doubling time were different between studies. In the Harrison study, the mortality doubling time was slightly increased by CR (p = 5.83*10^-4^), while in the Svenson study there was no significant effect on the mortality doubling time (p = 0.16). The effect of CR on baseline hazard also differed by study: in the Harrison study, mice experiencing 30% CR had a lower baseline hazard (p = 3.83*10^-9^), but there was little effect of CR on baseline hazard in the Svenson study (p = 0.11). These results also contrasted from previously reported effects of 20% and 40% caloric restriction in the DRiDO study, in which caloric restriction significantly reduced the mortality doubling time with no significant effect on baseline hazard (**Figure 1F**) (Di Francesco et al. 2023).

The effects of rapamycin treatment on aging rate and baseline hazard were also inconsistent across study (**Figure 1F**). Despite increasing lifespan in both studies, rapamycin treatment did not significantly impact baseline hazard (p = 0.63) or mortality doubling time (p = 0.20) in the Harrison study. In contrast, in the Svenson study, rapamycin treatment significantly decreased the mortality doubling time(p = 1.33*10^-10^) while also significantly increasing baseline hazard (p = 7.49*10^-7^).

In the Shock study, neither baseline hazard (p = 0.075) nor demographic aging rate (p = 0.31) significantly varied with sex.

### Study-specific genome-wide association analyses revealed five non-overlapping lifespan QTL

To investigate the genetic basis of lifespan, 1009, 460, and 916 mice were genotyped from the Harrison, Shock, and DRiDO studies, respectively (**Data S2 - S7**; Svenson study mice were not genotyped, see **METHODS**). We used residual maximum likelihood (REML) (Broman et al. 2019) to estimate the narrow-sense heritability (*h*^2^) of lifespan in each study after correcting for sex (Shock) or intervention (Harrison and DRiDO studies), as well as DO generation, which may impact allele frequencies and recombination density. Genetics explained 18.5% (SE = 6.5%) of variance in lifespan in the Harrison study, 15.0% (SE = 15.0%) in the Shock study, and 24.6% (SE = 7.7%) in the DRiDO study.

Genome-wide scans for quantitative trait loci (QTL) performed separately on lifespans from each of the Harrison, Shock, and DRiDO studies identified three unique QTL at a permutation-based significance threshold (**Figure 2A-C**; ɑ = 0.05, 1000 permutations; **Table S2**) (Doerge and Churchill 1996a). Two of these were identified in only one study: chromosome 12 at 82.92 Mb (2LOD support interval, or ‘2LOD SI’ = 81.29-83.37; DRiDO study), and chromosome 16 at 7.11 Mb (2LOD SI = 5.84-8.07; Shock study). Overlapping loci on chromosome 18 were identified from the DRiDO study (21.68 Mb; 2LOD SI = 20.46-25.82) and Harrison study (21.67 Mb; 2LOD SI = 15.81-25.76). At a lower, but still conservative, significance threshold (LOD ≥ 6; see **METHODS**), two additional QTL were identified from the DRiDO study (**Figure 2C**): both on chromosome 7, at 11.39 Mb (2LOD SI = 6.76-12.71) and 108.57 Mb (2LOD SI = 101.3-142.98) (**Figure S1**).

**Figure 2.**
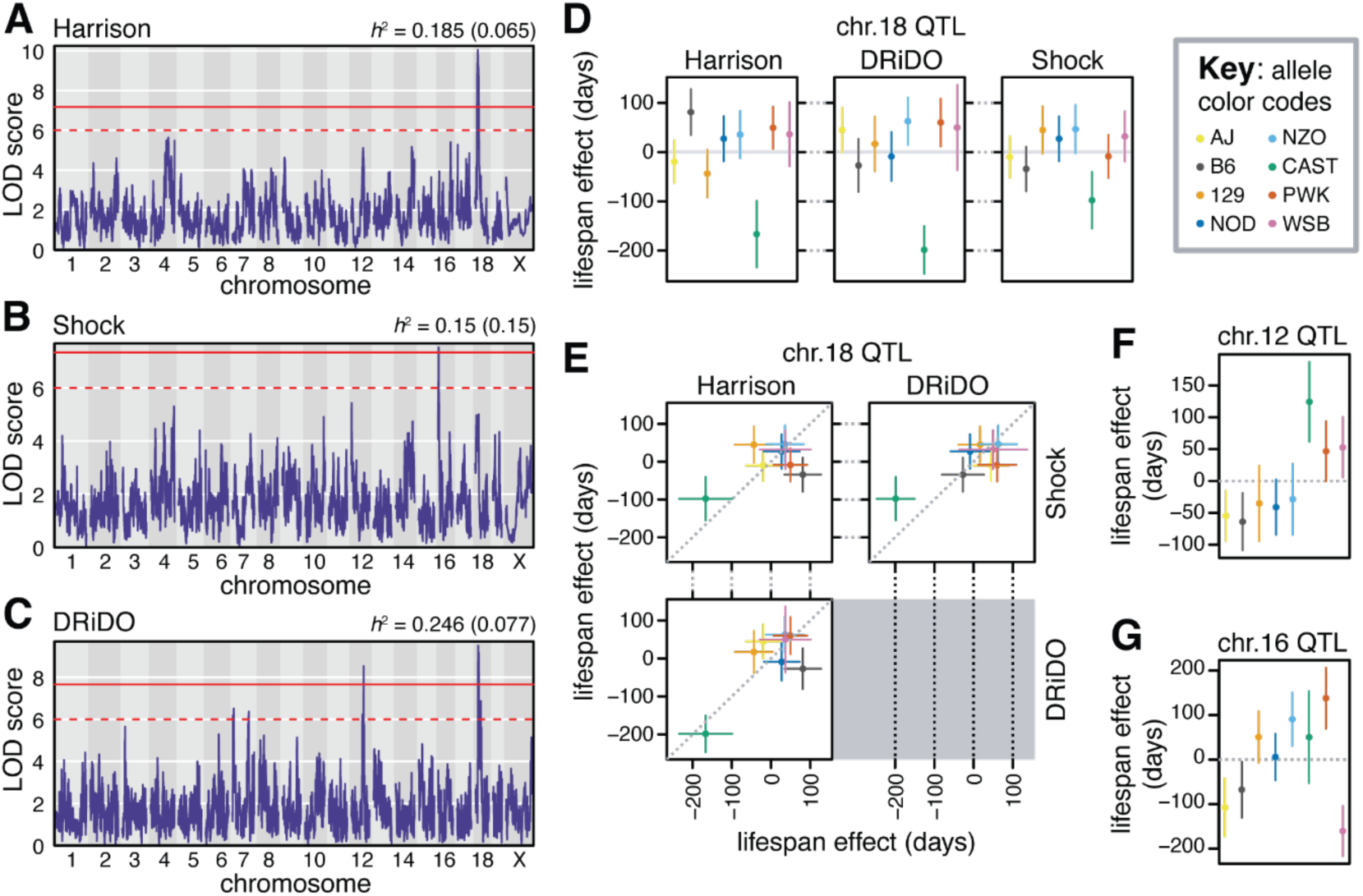
Genetic analysis of lifespan in DO mice. **A,** Manhattan plot of the additive genome-wide scan on lifespan in the Harrison study. The solid red line indicates genome-wide significance, while the dashed red line indicates a significance threshold of LOD 6. The narrow-sense heritability (h^2^) of lifespan is reported, with standard error about the h^2^ estimate reported in parentheses (*top right*). **B,** Manhattan plot of lifespan in the Shock study. **C,** Manhattan plot of lifespan in the DRiDO study. **D,** Allelic effects (BLUPs) and corresponding standard errors of the chromosome 18 locus detected in the Harrison (*left*), DRiDO (*middle*), or Shock (*left*) study. **E,** Comparisons of allelic effects (BLUPs) and standard errors of the chromosome 18 locus in each pair of studies. **F,** Allelic effects (BLUPs) and corresponding standard errors of the chromosome 12 locus detected in the DRiDO study. **G,** Allelic effects (BLUPs) and corresponding standard errors of the chromosome 16 locus detected in the DRiDO study.

For each QTL, the allelic effect of each founder haplotype on lifespan was estimated. The non-overlapping QTL on chromosome 18 had the largest effects on lifespan, explaining 4.7% of phenotypic variance in the DRiDO study and 4.5% of variance in the Harrison study. The effects of these loci were primarily driven by negative effects of the CAST/EiJ (CAST) allele. Animals with one or more CAST alleles at this locus lived 83.4 fewer days on average in the Harrison study (9.0%) and 99 fewer days on average in the DRiDO study (11.8%, **Figure 2D**). We presume these to represent the same true QTL, as the effects of each founder haplotype were significantly correlated in each study (Pearson correlation = 0.763, p = 0.027, **Figure 2E**). In the Shock study, this locus failed to reach genome-wide statistical significance; the effects of the founder haplotypes were significantly correlated with the DRiDO study (Pearson correlation = 0.820, p = 0.013) but not the Harrison study (Pearson correlation = 0.557, p = 0.151) and the effect of the CAST allele on lifespan at this site was approximately half as large as in the DRiDO or Harrison studies (**Figure 2D-E**).

The QTL on chromosomes 12 and 16 explained 4.2% and 6.9% of variance in lifespan within the DRiDO and Shock studies, respectively. Mice containing one or more CAST alleles at the chromosome 12 locus lived 62.1 days longer (7.4%) on average in the DRiDO study (**Figure 2F**). Notably, this locus was not detected at genome-wide significance when excluding mice that did not survive until the initiation of dietary intervention (6 months). In the Shock study individuals with at least one Watkins Star Line B (WSB) allele at the chromosome 16 locus lived 80.1 fewer days (10.8%) on average, and individuals with at least one PWK allele lived 69 days longer on average (9.3%, **Figure 2G**).

For each study, we divided the total fraction of phenotypic variance explained by the detected QTL by the variance explained by genetics overall (i.e. the heritability) to infer the mapped percentage of heritable variation. Considering only QTL significant at a genome-wide threshold, 24.3% (0.045/0.185), 46.0% (0.069/0.150), and 36.2% (0.089/0.246) of heritable variation in lifespan was explained in the Harrison, Shock, and DRiDO studies, respectively. Inclusion of the two QTL in the DRiDO study on chromosome 7, which each explained 3.2% of the total variance in lifespan in the DRiDO study, increased the percentage of explained heritable variation in this cohort to 62.2% (0.153/0.246). The portion of lifespan variances attributable to genetics but remaining unmapped were 14.0%, 8.1%, and 9.3% in the Harrison, Shock, and DRiDO studies, respectively.

### Meta-analysis of DO lifespan identifies three additional QTL

To improve power to detect QTL associated with longevity in DO mice, we performed a meta-analysis of lifespan using combined data from the 2,395 mice in the Harrison, Shock, and DRiDO studies (**Data S8 - S9**). After correcting for study, sex, intervention and generation wave, *h*^2^ of combined lifespan measurements across the three cohorts was 18.6% (SE = 3.8%). A whole-genome scan identified two previously identified QTL, chromosome 12 position 81.19 (2LOD SI = 74.9-84.64) and chromosome 18 at21.81 Mb (2LOD SI = 20.29-25.38) at genome-wide significance (**Figure 3A; Table S2**). The chromosome 16 locus, previously identified in the Shock cohort, was also detected at a significance level of LOD ≥ 6 at 8.82 Mb (2LOD SI = 6.46-10.46).

**Figure 3.**
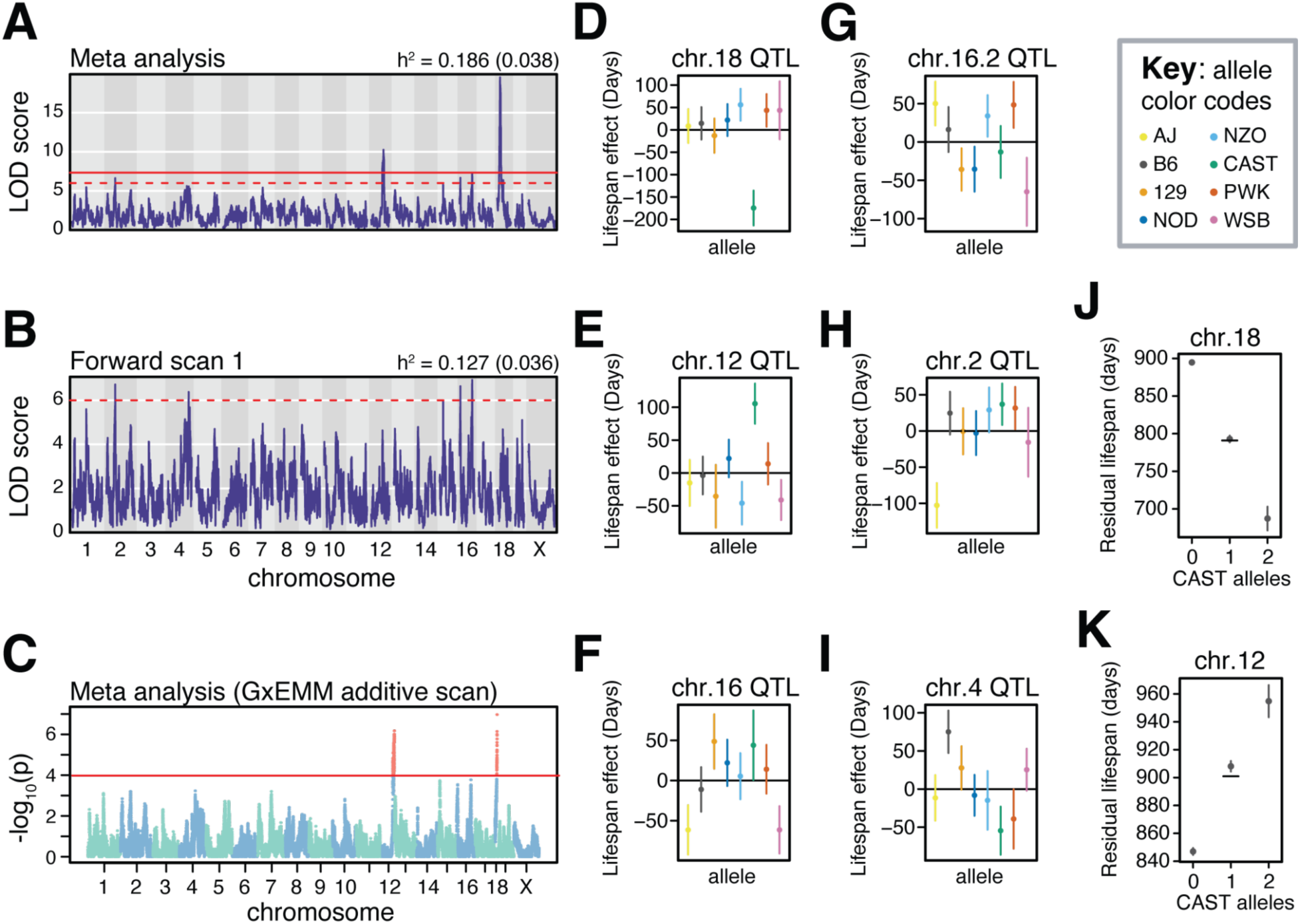
Meta analysis of lifespan in DO mice. **A,** Manhattan plot of the additive genome-wide scan on lifespan using combined data from the Harrison, Shock, and DRiDO studies. The solid red line indicates genome-wide significance, while the dashed red line indicates a significance threshold of LOD ≥ 6. The narrow-sense heritability (h^2^) of lifespan is reported, with standard error about the h^2^ estimate reported in parentheses (*top right*). **B,** Manhattan plot of the forward regression scan on the combined data, in which statistically significant loci were included as additive covariates in the model. **C,** Manhattan plot of the additive genome-wide scan on combined lifespan data using GxEMM. Markers reaching statistical significance are colored red. **D,** Allelic effects (BLUPs) and corresponding standard errors of the chromosome 18 locus previously detected in the Harrison and DRiDO studies. **E,** Allelic effects (BLUPs) and corresponding standard errors of the chromosome 12 locus previously detected in the DRiDO study. **F,** Allelic effects (BLUPs) and corresponding standard errors of the chromosome 16 locus previously detected in the Shock study. **G,** Allelic effects (BLUPs) and corresponding standard errors of the second chromosome 16 locus. **H,** Allelic effects (BLUPs) and corresponding standard errors of the chromosome 2 locus. **I,** Allelic effects (BLUPs) and corresponding standard errors of the second chromosome 4 locus detected in the forward regression. **J,** Mean residual lifespan and standard error of DO mice as a function the number of *CAST* alleles at the chromosome 18 locus. The black line denotes the expected residual lifespan of individuals carrying a single *CAST* allele at this locus. **K,** Mean residual lifespan and standard error of DO mice as a function the number of *CAST* alleles at the chromosome 12 locus.

While neither of the QTL on chromosome 7 were reproducibly detected in this meta analysis, novel loci on chromosomes 2 (59.34 Mb; 2LOD SI = 11.04-159.99) and 16 (83.26 Mb; 2LOD SI = 79.84-84.56) were detected at our lower significance threshold. Additionally, a forward scan was performed in which each of the genome-wide significant loci were added to the mixed effects model as additive covariates (**Figure 3B**). The forward scan identified an additional locus on chromosome 4 at 137.61 Mb (2LOD SI = 88.47-147.61). The loci on chromosomes 18 and 12 were reproducibly detected using an independent method, GxEMM (Wright et al. 2022), at a previously established significance threshold of p ≤ 10^-4^ (**Figure 3C**).

The effects of chromosome 18 and 12 appeared to be driven by the same alleles detected using data in the DRiDO study, with the CAST allele associated with a decrease in lifespan of 175 days at chromosome 18 and an increase in lifespan of 106 days at the chromosome 12 locus (**Figure 3D-E**). Additional QTL displayed more complex patterns of allelic effects, with two or more alleles influencing lifespan at these sites (**Figure 3F-I**). Mice carrying at least one copy of CAST at the chromosome 18 site had significantly decreased lifespans (log-rank test, ꭓ^2^ = 40.53, p = 1.58*10^-9^) and mice carrying at least a single copy of CAST at the chromosome 12 locus had significantly increased lifespans (log-rank test, ꭓ^2^ = 22.36, p = 1.39*10^-05^). After correcting for study, diet, sex, and DO generation, the effect of both loci on lifespan is additive (**Figure3J-K**).

Across cohorts, the QTL on chromosome 18 explained 3.7% of phenotypic variance in lifespan and the QTL on chromosome 12 explained 2% of phenotypic variance. Collectively, the four QTL on chromosomes 2, 4 and 16 explained 5.2% of variance in lifespan. Together, loci detected in the meta analysis explained 10.9% of overall trait variance and 58.6% of the estimated heritable variance for lifespan. Assuming that remaining additive effects explain less phenotypic variance than the QTL with the smallest effect (1.2%), there would need to be at least 7 undiscovered QTL influencing lifespan in this combined cohort in order to account for the remaining 7.7% of total variance that was heritable but remained unexplained.

### Association mapping of genes underlying lifespan QTL

In order to localize the effects of lifespan QTL, we performed variant association mapping of loci detected at the permutation-based significance threshold (**Figure 4**) or at LOD ≥ 6 (**Figure S2**) in the meta-analysis (**Table S3**). The 2 LOD support intervals around the chromosome 18 loci in the Harrison and DRiDO studies encompassed ∼26 protein coding genes (**Table S3**). Variant association mapping further refined the effect of this locus to 10 genes: B4galt6, Ttr, Trappc8, Rnf125, Rnf138, Mep1b, Garem1, Klhl14, Ccdc178, and Axl3 (**Figure 4A**). Association mapping of the chromosome 12 locus localized its effect to a region encompassing ∼18 genes, including DRAGO/*Susd6* (Polato et al. 2014), a DNA damage-responsive tumor suppressor and *Rsg6*, a regulator of G-protein signaling (**Figure 4B**). Association mapping of the chromosome 16 locus localized its effect to ∼8 genes: Rbfox1, Tmm114, Mettl22, Tmem186, Pmm2, Abat, Carhsp1, and Usp7 (**Figure 4C**).

**Figure 4.**
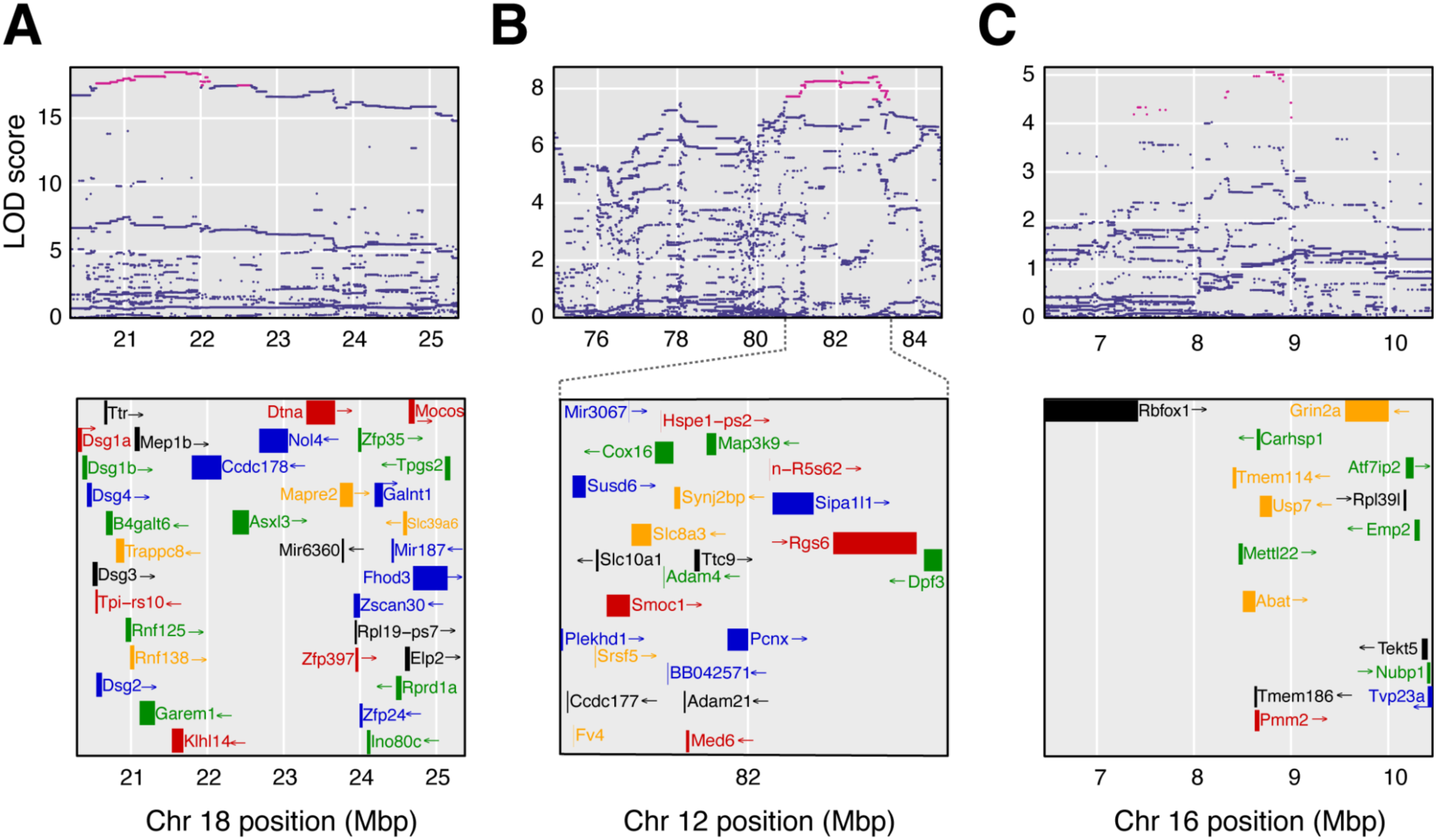
Variant association mapping of loci detected at statistical significance in one or more studies. **A,** Variant association mapping of the QTL on chromosome 18 depicting the LOD scores for each variant within the 3 LOD support interval around the locus in the meta-analysis (*top*). The area encompassed by the plot corresponds to the 3 LOD support interval around the locus in the meta-analysis. The most likely candidate SNPs are highlighted in pink. Genes within the genomic interval are depicted (*bottom*). **B,** Variant association mapping of the chromosome 12 locus in the meta-analysis. Here, the *bottom* panel is zoomed in on the coordinates of the most likely causal variants for clarity (dashed lines). **C,** Variant association mapping of the chromosome 16.1 locus in the meta-analysis.

### Loci interact with sex and dietary restriction to influence lifespan in DO mice

Four of the five unique QTL identified in the individual studies only emerged in a single study. We evaluated the potential relevance of these five QTL across studies by fitting single-QTL models at the peak QTL positions using data from their respective studies, including interaction terms for sex (Shock study) or dietary intervention (DRiDO study). While none of the QTL from the DRiDO study were found to have interactions with dietary intervention, the locus on chromosome 16 had a sex-specific effect on lifespan in the Shock cohort (single locus ɑ = 0.05, 1000 permutations; **Figure 5A**; **Table S4**). The largest effects on lifespan at this locus were observed in female animals: after correcting for sex and DO generation, female mice carrying at least one copy of the PWK allele lived 150 days (SE = 18.73) longer than males, with the allele having a minimal effect on lifespan in males. None of the QTL were found to have interactions with sex or dietary intervention in the meta analysis (ɑ = 0.05, 1000 permutations).

**Figure 5.**
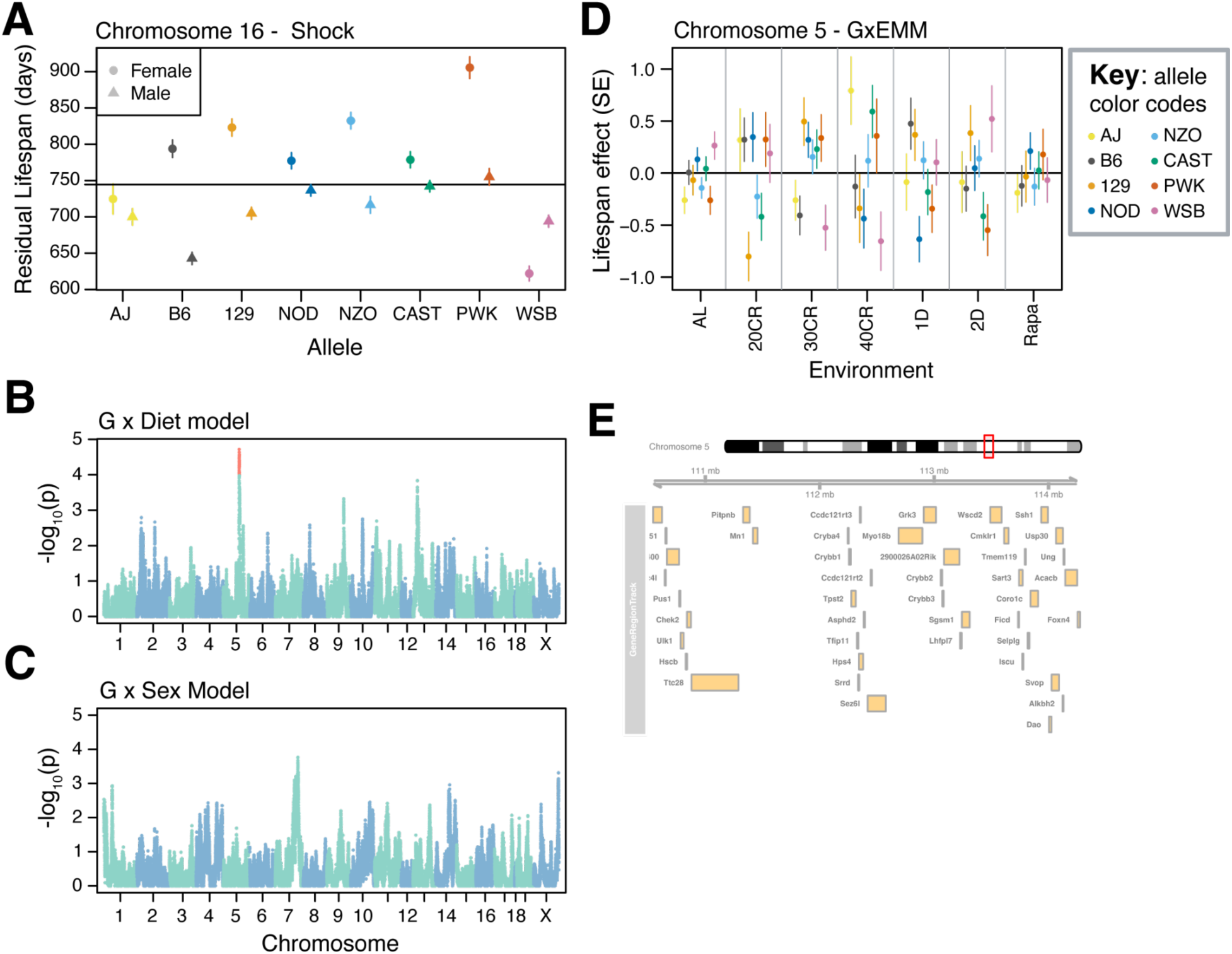
Genome-wide scans for loci with diet or sex specific effects on lifespan. **A,** Sex-specific effects of the chromosome 16 locus within the Shock study. Mean lifespan after correcting for sex and generation wave is shown by the black line. Error bars denote standard error. **B,** Whole genome scan for loci influencing lifespan via interaction with diet using GxEMM. Red points denote loci with significant effects at p < 1*10^-4^. **C,** Whole genome scan for loci influencing lifespan via interaction with sex using GxEMM. **D,** Allelic effects of the chromosome 5 locus on lifespan as a function of dietary intervention or drug treatment, with standard errors. **E**, Map of the 2LOD support interval about the peak marker at chromosome 5 depicting all protein coding genes within the interval.

Motivated by the identification of a sex-responsive lifespan QTL, we conducted whole-genome meta analyses using gene-by-environment mixed models (GxEMM) to detect loci with treatment and/or sex dependent effects on lifespan (Wright et al. 2022). Whole-genome scans for loci with diet-specific or sex-specific effects on lifespan resulted in detection of a single diet-responsive locus on Chromosome 5 at 112.74 Mb (2LOD SI= 108.70, 115.39, -log_10_p > 4, **Figure 5B**). No sex-responsive loci were detected at a genome-wide significance threshold (**Figure5C**). This locus exhibited the strongest effects on lifespan under 40% CR, with the largest effects driven by alleles from the AJ and WSB founding strains (**Figure 5D**). The 2LOD support interval about this locus spanned 3.78 megabases and contained 44 protein coding genes, limiting our ability to identify an obvious candidate (**Figure 5E, Table S3**).

## DISCUSSION

Motivation for the initiation of these studies was to evaluate the effects of dietary interventions, rapamycin treatment, and sex on lifespan in the genetically diverse background of DO mice. In addition to differences in cohorts, facilities, and personnel, the studies varied in their experimental designs, especially with respect to the additional phenotyping, feeding schedules, diet composition, housing temperature, and co-housing of the animals (Luciano and Churchill 2024; Di Francesco et al. 2023). For instance, mice in the Harrison study were fed non-irradiated 5LG6 chow, which differs in both micronutrient composition and fat content from the 5K52 chow used in the other studies, and could result in different effects of interventions on lifespan. Despite these differences, caloric restriction and rapamycin treatment robustly increased lifespan across all studies (**Figure 1**). Of the interventions analyzed, CR had the greatest effect on lifespan, with intermittent fasting and rapamycin treatment also significantly increasing lifespan but to a lesser degree. While the lifespan-extending effects of these interventions were consistent across studies, their specific effects on the shapes of the survival curves (baseline hazard and/or mortality doubling time) were less consistent, highlighting the difficulty of modeling Gompertzian mortality parameters.

While the heritability and genetics of lifespan varied from study to study, a locus on chromosome 18 was detected in two independent studies of DO lifespan (Harrison and DRiDO), suggesting it to be a significant contributor to longevity in DO mice. A second locus on chromosome 12 was detected in the DRiDO study, and both loci were detected at a higher level of significance in meta-analysis across studies, implying their broader relevance as lifespan QTL. While causes of mortality were not explored in any study, the detection of QTL unique to individual studies raises the possibility that differences in experimental design, can influence the genetics of lifespan, as different causes of mortality may be more important under different experimental conditions; for example, different dietary interventions have been shown to have contrasting effects on health-related phenotypes.

Although CR and rapamycin treatment consistently increased lifespan across genetically distinct individuals and independent studies, the detection of a novel QTL with intervention-specific effects on lifespan demonstrates that these interventions may not have the same effects across all individuals. This QTL had the largest effect under various dietary interventions and little effect on longevity in rapamycin-treated mice. Given that previously identified QTL have been shown to impact lifespan through body weight (Bou Sleiman et al. 2022), it is possible that the effect of this locus is also mediated by body weight, and only under appropriate dietary conditions. Diet-specific effects of loci may explain the contrasting effects of dietary intervention across health-related traits, or why certain individuals appear to benefit more from particular interventions. The extent to which interactions between genes and the environment influence lifespan in DO mice remains unclear but warrants further exploration.

Aggregation of multiple datasets empowered the detection of several novel lifespan QTL, comparable to other large-scale studies of lifespan in outbred populations such as UM-HET3 (Miller et al. 2007; Bou Sleiman et al. 2022). Meta analysis revealed six lifespan QTL, including all three of the loci detected at genome-wide significance in individual studies. These results, combined with our estimate of heritability, allowed us to roughly estimate that at least 12 QTL would be required to fully explain the heritable variation in lifespan in the meta-population. Our results provide evidence for a complex, polygenic genetic architecture underlying lifespan in DO mice.

## METHODS

### Study Designs

#### Dietary Restriction (DRiDO)

This study is extensively described elsewhere (Di Francesco et al. 2023). Briefly, female DO mice were received at ∼4 weeks of age in 12 waves from March 2016 through November 2017. Mice were housed in groups of 8 in single large-format ventilated pens with nestlets, biotubes, and gnawing blocks. Mice were randomized to one of five dietary interventions which were initiated for the surviving mice at 6 months of age: ad libitum (AL; n = 188), 1 day per week fasting (1D; n = 188), 2 days per week fasting (2D; n = 190), 20% caloric restriction (20; 2.75g/mouse/day; n = 189), and 40% caloric restriction (40; 2.06g/mouse/day; n = 182). Mice were extensively phenotyped as described (Di Francesco et al. 2023) and maintained until they died naturally. The mouse room was on a 12/12 hour light/dark schedule from 6:00 am to 6:00 pm and kept at 73° +/- 2° F.

#### Harrison

Founder DO mice (167 retired breeder pairs) were obtained from the Jackson Laboratory and female offspring were accumulated for the lifespan study over 5 months. All mice were microchipped at 4 weeks of age. Mice were housed 22 per large-format double pens connected by a tunnel on an open-air rack. All 22 mice per pen were from different breeder pairs. Pens had pine shaving bedding with acidified water. Every week, one pen of the connected pair would be changed. Mice would be herded into one pen and the tunnel blocked off. The dirty pen would be removed, and a new clean pen attached with fresh water and grain. The ad libitum control mice (n = 349) were on a non-irradiated diet (5LG6, TestDiet, Purina) from weaning. The diet restricted (n = 335) mice received 2.2 g/day/mouse of ground non-irradiated diet via modified fish feeders that were programmed to dump the ground diet onto the floor of the cage between 6-7 pm after lights were off. Modified feeders were restocked every 7 days. Any grain left in the feeders after 7 days was dumped on the cage floor. Proper feeder performance was indicated by a weighted string that was wound around a screw when the feeders dumped food. Diet restriction began at 4 weeks of age after being microchipped. An additional 339 mice received non-irradiated diet until they were 16 months of age, whereupon they started on 5LG6 diet with 142 ppm encapsulated rapamycin (Rapamycin Holdings, actual concentration of rapamycin in diet is 14 ppm, TestDiet, Purina). Mice were maintained until they died naturally. The mouse room was on a 12/12 hour light/dark schedule from 6:00 am to 6:00 pm and kept at 73° +/- 2° F.

#### Svenson

We obtained female DO mice from the Jackson Laboratory breeding colony at ∼4 weeks of age. Mice were obtained in eight waves over the course of 1 year and enrolled by randomization to dietary intervention protocols. Mice were housed 8 per group in single large format pens. Interventions were implemented as described for the Harrison study with a few differences indicated here. The ad libitum fed control mice (n = 319) were on a 4% irradiated diet (5K52, TestDiet, Purina). The diet restricted mice (n = 316) received 2.2 g/day/mouse of ground 4% irradiated diet. DR mice were fed at ∼7am daily and food was placed directly onto the bottom of the pen by a technician. On Friday the DR mice received a triple feeding (6.6 g/mouse) and were fed again on Monday morning. A third group of mice (n = 317) were maintained on the ad libitum protocol until 16 months of age and were then switched to rapamycin diet as described above. Mice experienced minimal handling (monthly body weights and weekly pen changes) and were maintained until they died naturally. The mouse room was on a 12/12 hour light/dark schedule from 6:00 am to 6:00 pm and kept at 70° +/- 2° F.

For this study, DNA samples were collected for genotyping on the MUGA array, as described for other studies below. However, irregularities with sample labeling/handling made us question the integrity of our ID matches between mice and samples. We performed quality-control assessment by comparing genotype-predicted versus recorded coat colors across the mice (Silvers 2012) and confirmed extensive sample mismatches (data not shown). We therefore excluded these data from our genetic analyses.

#### Shock

We obtained female (n = 244) and male (n = 240) DO mice from the Jackson Laboratory breeding colony at ∼4 weeks of age. Mice were obtained in five waves from June 2011 through August 2012. Mice were housed in single-sex groups of 5 in standard ventilated duplex pens. All mice were fed ad libitum on 6% sterilized gain (5K52, TestDiet, Purina). Mice experienced minimal handling (body weights and other non-invasive procedures). At 6, 12, and 18 months we obtained 3x100ul retroorbital blood draws, with 2 weeks recovery time between each. Mice were maintained until they died naturally. The mouse room was on a 12/12 hour light/dark schedule from 6:00 am to 6:00 pm and kept at 70° +/- 2° F.

All procedures used in these studies were reviewed and approved by the Jackson Laboratory Animal Care and Use Committee.

### Genotyping

Genotypes for all studies reported here were obtained using the mouse universal genotyping array (MUGA) (Morgan et al. 2016). DNA was isolated from tail tips using standard methods and shipped to Neogen Genomics (Lincoln, NE, USA) for analysis. Samples were genotyped using the MUGA (Harrison), MegaMUGA (Shock), or GigaMUGA (DRiDO) genotyping arrays. Founder haplotypes were reconstructed using the R/qtl2 software and samples with call rates at or above 90% were retained for analysis. Genome coordinates were from mouse genome GCRm39 and gene locations were taken from the Mouse Genome Informatics databases (Blake et al. 2021).

### Data analysis

#### Survival analysis

We compared survival among each cohort of animals, as well as between experimental groups within studies. This was done by plotting Kaplan-Meier curves and by testing the equivalence of survival distributions among each cohort or experimental group using log-rank tests using overall tests (across cohorts and within each cohort) as well as pairwise comparisons between each experimental group and its respective within-study control group (ex: comparing rapamycin treatment to *ad libitum* within the Harrison study). p-values are reported with no correction for multiple comparisons and are considered significant at p < 0.05. Median lifespan was estimated in each cohort as well as within each experimental group. The effects of dietary interventions and/or sex were estimated via Cox proportional hazards regression analysis and are reported as hazard ratios with 95% confidence intervals. p-values are reported without correction for multiple comparisons and are considered significant at p < 0.05. Survival analysis was conducted using the “survival” (Therneau and Grambsch 2000; Therneau 2024) package in R and plotted via the “ggsurvfit” package (Sjoberg et al. 2024). Mortality doubling times and baseline hazards were estimated, beginning at the time of intervention, from a Gompertz log-linear hazard model with a 95% confidence interval and percentage change relative to female mice on an *ad libitum* diet via the “flexsurv” package in R (Jackson 2016).

#### Additive whole-genome scans

All genetic analysis was conducted using the “rqtl2” package in R (R Core Team 2024; Broman et al. 2019). Whole-genome scans for lifespan QTL were carried out via the scan1() function using a mixed effects model in which lifespan was regressed on 8-state allele probabilities for each individual in a dataset and LOD scores were recorded at each marker (Broman and Sen 2009). Within the Dietary Restriction and Harrison studies, dietary intervention and DO generation were included as additive covariates. In the Shock study, Sex and DO generation were included as additive covariates. In the meta-analysis, Study, Diet, Sex, and DO generation were included as additive covariates. For each genome-wide scan, 1000 permutations of the data were performed in which phenotypes were randomized and a whole-genome scan was run (Doerge and Churchill 1996b). The maximum LOD score observed in each permuted scan was recorded, and the 95^th^ percentile of the distribution of 1000 maximum LOD scores was used as the significance threshold (ɑ = 0.05). QTL with LOD scores greater than this threshold were considered significant at our permutation-based threshold. In addition to this significance level, a significance level of LOD ≥ 6 was also used to identify loci contributing to variance in lifespan. While less conservative, this threshold is more stringent than a previously reported method (Wright et al. 2022). We report 2 LOD support intervals (‘2LOD SI’), corresponding to a 2 LOD drop around each peak position, about each peak marker identified in whole-genome scans.

#### Forward regression analysis

In the meta-analysis, forward regression analysis was performed to account for the effects of genome-wide significant QTL when searching for additional loci influencing lifespan. This was done via the scan1() function using a mixed effects model including dietary intervention, sex, and DO generation as additive effects and kinship as a random effect. In addition to these covariates, previous QTL identified at a genome-wide significance level were included in the model as additive effects. QTL were encoded as numeric variables representing the genotype state at the marker with the highest LOD score as reported by association mapping. Only QTL reaching permutation-based significance thresholds were included in the model as additive covariates.

#### Effect size estimation and percent variance explained

Best linear unbiased predictors (BLUPs) and corresponding standard errors were computed for all QTL using the scan1blups() function in “rqtl2” using the additive covariates listed above. Phenotypic variance explained by each QTL was calculated using the following formula (Broman and Sen 2009):

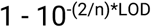

Where n is the number of samples in a particular dataset and LOD corresponds to the LOD score of the peak marker at each QTL.

#### Variant association

*<u>Variant association mapping</u>* was conducted within a 2LOD drop of the peak position associated with each QTL via the scan1snps() function in “rqtl2” using the same additive covariates listed in the *Additive whole-genome scans* section above. Variant and gene SQLite datasets used in this analysis are available at the “rqtl2” user guide website: https://kbroman.org/qtl2/assets/vignettes/user_guide.html.

#### Single-QTL models for diet- and sex-specific loci

Within individual studies, QTL were tested for interaction with experimental factors unique to those studies. In the Shock cohort, QTL were tested for interaction with sex, while in other cohorts QTL were tested for interactions with one or more of the dietary interventions. Interaction tests were not conducted genome-wide using “rqtl2”, since this package does not fit interactions as random effects. For each QTL, tests were run using the 8-state allele probabilities at peak positions identified in whole-genome scans using the fit1() function in “rqtl2”. To assess significance, two models were run: an additive model in which experimental factors (sex, dietary intervention) and DO generation were included as additive covariates and an interaction model that included the same additive covariates with an additional interaction term corresponding to the experimental factor being tested. The reported LOD is the LOD of the interaction model minus the LOD of the additive model. To establish significance, 1000 permutations were run at each QTL in which phenotypes were randomized before running the additive and interaction models. After ordering the 1000 resulting LOD scores, the 95^th^ percentile was chosen as the significance threshold (ɑ = 0.05). Interactions between QTL and experimental conditions were considered significant if their LOD was greater than the ɑ = 0.05 threshold. Interaction effects were plotted as residual lifespan values as a function of sex after correcting for additive covariates.

#### Genome-wide scans for diet- and sex-specific loci

Gene by environment mixed effects models (GxEMM) were run using the ‘do-qtl’ software package in Python version 3.8.16 (Wright et al. 2022). Study, sex, and diet were included in the model as fixed effects, while diet and kinship were supplied as random effects. Error estimates were computed based on 1,000 permutations of the data. A significance threshold of p <= 1*10^-4^ was used to define loci as statistically significant (Wright et al. 2022).

## Supplementary Figures

**Figure S1.**
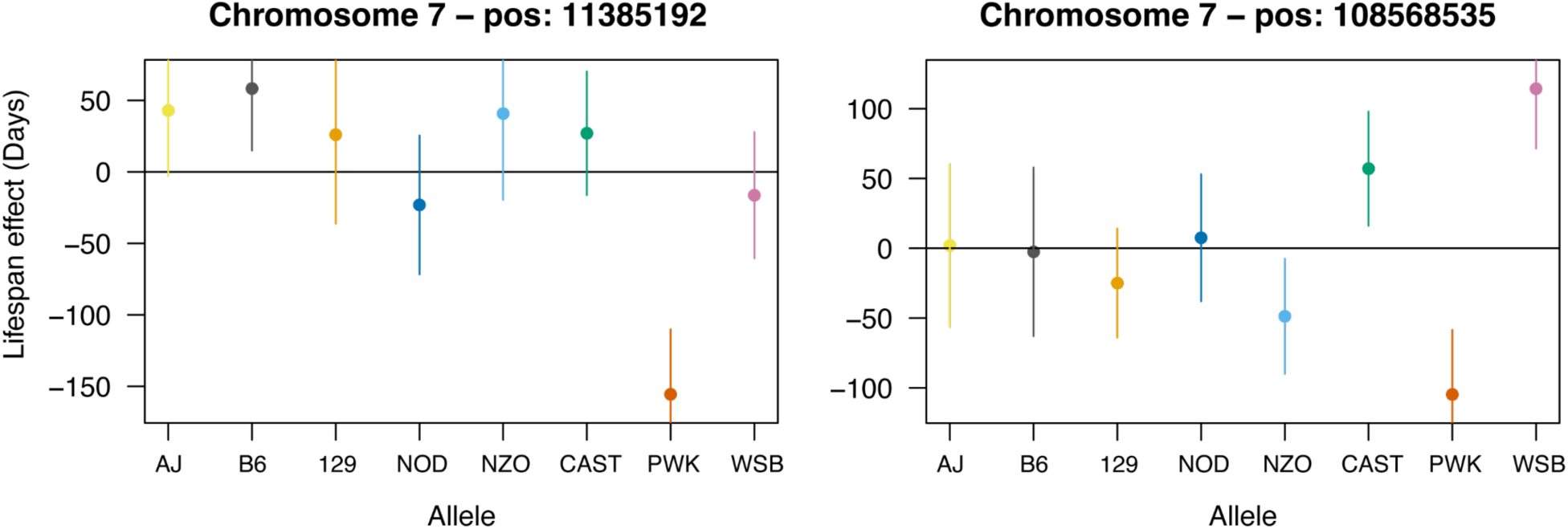
Effects of peak markers at additional QTLi (DRiDO study) on lifespan. Effects (BLUPs) and corresponding standard errors for animals containing at least one copy of the corresponding founder allele.

**Figure S2.**
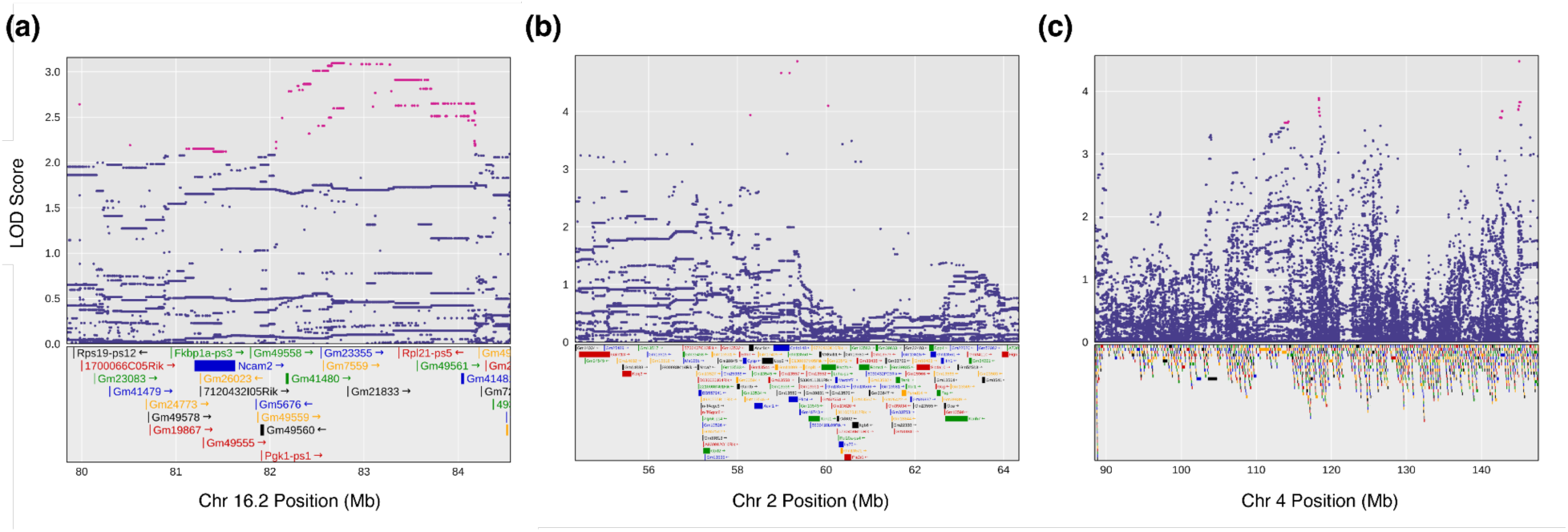
Association mapping of nominally significant QTL from a meta-analysis of lifespan. **A,** Variant association mapping of the QTL on chromosome 16.2 depicting the LOD scores for each variant within the 3 LOD support interval around the locus in the meta-analysis (*top*). The area encompassed by the plot corresponds to the 3 LOD support interval around the locus in the meta-analysis. The most likely candidate SNPs are highlighted in pink. Genes within the genomic interval are depicted (*bottom*). **b,** Variant association mapping of the chromosome 2 locus in the meta-analysis. **c,** Variant association mapping of the chromosome 4 locus in the meta-analysis.

## Supplementary Tables / Data (available at Dryad link below)

### Data S1

Lifespan and covariate data for the mice in each of the four studies. ‘Study’ corresponds to the study the animal was enrolled in: ‘Shock’, ‘Harrison’, ‘Svenson’, or ‘DRiDO’ (Dietary Restriction). ‘Mouse.ID’ corresponds to the unique identifier assigned to each animal. ‘Sex’, ‘Diet’, and ‘Generation’ are the primary covariates used in the analysis; ‘Generation’ corresponds to the specific cohort or generation wave of the DO mice. ‘Status’ reflects the outcome of animals as they exit their respective study; ‘1’ means the animal died and ‘0’ means the animal was censored.

### Data S2

Eight-state allele probabilities from the Harrison study.

### Data S3

Lifespan and covariate data for the Harrison study.

### Data S4

Eight-state allele probabilities from the Shock study.

### Data S5

Lifespan and covariate data for the Shock study.

### Data S6

Eight-state allele probabilities from the Dietary Restriction (‘DRiDO’) study.

### Data S7

Lifespan and covariate data for the Dietary Restriction (‘DRiDO’) study.

### Data S8

Combined eight-state allele probabilities from the Harrison, Shock, and DRiDO studies.

### Data S9

Combined lifespan and covariate data for the Harrison, Shock, and DRiDO studies.

### Table S1

Test statistics for log rank tests comparing lifespan among female mice fed an *ad libitum* diet by study. ‘Test’ refers to the pair of studies being tested. ‘chi_sq’ and ‘pval’ columns provide the ꭓ^2^ and p-value associated with each test.

### Table S2

List of QTL detected in the Harrison, Shock, and DRiDO studies, the meta analysis, and the GxEMM scans. ‘Study’ refers to the particular dataset in which a QTL was detected. ‘chr’ refers to the chromosome on which the QTL is located. ‘pos’ refers to the physical location on the respective chromosome at which the QTL is located in megabases (Mb). ‘lod’ refers to the LOD score of the peak marker at the QTL. ‘ci_lo’ refers to the location of the marker denoting the 2LOD drop prior to the peak marker. ‘ci_hi’ refers to the location of the marker denoting the 2LOD after the peak marker. ‘threshold’ refers to the significance threshold used to call the peak marker of the QTL, which can be ‘Nominal’, ‘Permutation’, or ‘GxEMM’. ‘Nominal’ corresponds to a conservative threshold of p<= 1*10^-6^, ‘Permutation’ corresponds to an alpha critical threshold of <=0.05 based on 1,000 permutations of the data, and ‘GxEMM’ corresponds to the previously reported threshold of p <= 1*10^-4^.

### Table S3

Table of genes underlying each QTL. ‘study’ corresponds to the study in which a particular QTL was mapped. ‘qtl_id’ corresponds to the chromosome and position at which the QTL was mapped, separated by a “_” character. ‘chr’ corresponds to the chromosome on which the QTL was mapped. ‘source’ corresponds to the database from which gene annotation data was collected. ‘type’ corresponds to the annotation associated with a marker (can be ‘gene’ or ‘pseudogene’). ‘start’ and ‘stop’ correspond to the first and last physical coordinates associated with the annotation in megabases (Mb), respectively. ‘strand’ corresponds to which strand of DNA the annotation is on. ‘ID’ corresponds to the database ID assigned to the annotation. ‘Name’ corresponds to the gene name associated with the annotation. ‘Dbxref’ lists annotation IDs in other databases. ‘gene_id’ corresponds to the gene ID in the corresponding database. ‘mgi_type’ corresponds to the gene type associated with the annotation in the corresponding database. ‘description’ provides a functional description of the protein or RNA associated with the annotation.

### Table S4

Statistical test for interaction between chromosome 16 and sex in the Shock study.

## Data Availability Statement

All supplementary Tables and Data, which includes primary genetic and phenotype data, are available at Dryad

[https://datadryad.org/stash/share/d5bGZs-WAhIFSHzywpRBIxJWN0mUFxysIe3zdbvmSwA]

## Acknowledgments

This work was funded by National Institutes of Health grants AG038070, AG022308, and AG079753, an Ellison Medical Foundation Senior Scholar Award (GAC), and by Calico Life Sciences LLC.

